# Neural responses to movie naturalistic stimuli are related to listening demand in cochlear implant users

**DOI:** 10.1101/2022.09.02.506424

**Authors:** Bowen Xiu, Brandon T. Paul, Joseph Chen, Trung Le, Vincent Lin, Andrew Dimitrijevic

## Abstract

There is a weak relationship between clinical and self-reported speech perception outcomes in cochlear implant (CI) listeners. Such poor correspondence may be due to differences in clinical and “real-world” listening environments and stimuli. Speech sounds in the real world are often accompanied by visual cues, background environmental noise and is generally in the context of a connected conversation. The aims of this study were to determine if brain responses to naturalistic speech could index speech perception and listening demand in CI users. Accordingly, we recorded high density EEG while CI users listened/watched a naturalistic stimulus (i.e., the television show, “The Office”). We used continuous EEG to quantify “speech neural tracking” (i.e., TRFs, temporal response functions) to the television show audio track and additionally 8–12 Hz (alpha) brain rhythms commonly related to listening effort. Background noise at three different signal-to-noise ratios (SNRs), +5, +10, and +15 dB were presented to vary the difficulty of following the television show mimicking a natural noisy environment. The task included an additional condition of audio-only (no video). After each condition, participants subjectively rated listening demand and the degree of words and conversations they felt they could understand. Fifteen CI users reported progressively higher degrees of listening demand and less words and conversation with increasing background noise. Listening demand and conversation understanding in the audio-only condition was comparable to that of the highest noise condition (+5 dB). The addition of the background noise reduced the degree of speech neural tracking. Mixed effect modeling showed that listening demand and conversation understanding were correlated to cortical speech tracking such that high demand and low conversation understanding lower associated with lower amplitude TRFs. In the high noise condition, greater listening demand was negatively correlated to parietal alpha power such that higher demand was related to lower alpha power. No significant correlations were observed between TRF/alpha and clinical speech perception scores. These results are similar to previous findings showing little relationship between speech perception and quality of life in CI users. However, the physiological responses to complex natural speech may anticipate aspects of quality-of-life measures such as self-perceived listening demand.

## 1 Introduction

Cochlear implants (CI) can successfully restore hearing for many individuals with profound-to-severe sensorineural hearing loss. Despite increases in post-implantation speech perception scores and quality-of-life (QoL) for most CI users (e.g. Dillon et al., 2018; Fabie et al., 2018), not all individuals achieve a favourable level of speech performance or QoL (e.g. Capretta & Moberly, 2016; Holden et al., 2013). Studies examining post-implantation outcomes have focused on etiology of hearing loss, duration of deafness, age at implantation, the amount of residual hearing, and device differences (e.g. Blamey et al., 2012; James et al., 2019; Kurz et al., 2019; Lazard et al., 2012). However, these factors only account for 10% to 22% of the outcome variability (Blamey et al., 2012; Lazard et al., 2012). Furthermore, the correlation between clinical speech perception test and subjective QoL questionnaire outcomes appears to be weak to moderate at best, with some studies reporting no significant relationship (e.g., McRackan et al., 2018; Ramakers et al., 2017; Thompson et al., 2020). Accordingly, QoL does not necessarily improve after cochlear implantation if a desired level of speech performance is reached. These observations motivate the development of clinical testing procedures that are sensitive to post-implant changes to QoL.

Clinical speech tests may fail to reliably predict QoL changes because current clinical testing materials do not fully capture aspects of real-world listening. Clinical tests take place with participants centered in a sound-attenuated booth with minimal reverberation where speech and background noise stimuli are presented at fixed, equidistant positions around the azimuth (Dunn et al., 2010; Minimum Speech Test Battery (MSTB) For Adult Cochlear Implant Users, 2011; Spahr et al., 2012). Additionally, the properties of speech and background noise stimuli themselves are often constant in terms of loudness and content. In contrast, listening in everyday life often occurs in dynamic complex environments where speech is often accompanied by fluctuating background noise, competing speech sources, and with varying access to visual (speech) input that influences speech comprehension. CI users often report that visual articulatory cues (i.e. lip, tongue, and teeth movements that accompany speech) are missing from clinical testing (Dorman et al., 2016). Visual articulatory cues are correlated with properties of the auditory speech signal and provide information that assists CI users in interpreting ambiguous speech, as demonstrated through the McGurk effect (Rouger et al., 2008; Stropahl et al., 2017). Results from neuroimaging studies suggest that visual articulatory cues increase selective attention, speech intelligibility, and neural speech tracking compared to audio-only speech-in-noise tasks, especially in individuals with hearing loss (Zion Golumbic et al., 2013a; O’Sullivan et al., 2019; Puschmann et al., 2019). On the other hand, background noise and reverberation negatively affects word recognition and listening effort (Picou et al., 2016). The quality of neural speech tracking has also been shown to decrease in the presence of noise or competing sound sources; background noise has been demonstrated to attenuate the brain response around 100 ms after a change in the speech envelope amplitude (Ding and Simon, 2012; Fuglsang et al., 2017; Petersen et al., 2017).

In these instances, the burden of degraded speech and environmental distractions can increase the overall cognitive demand required to adequately comprehend speech (Rönnberg et al., 2010; Picou et al., 2016). Listening to speech in noise can impose a strong demand on cognitive resources due to increased engagement of working memory (Rönnberg et al., 2010, 2013) and attention processes (Shinn-Cunningham and Best, 2008). In complex listening environments where the sound quality of speech is reduced, listeners must apportion their limited cognitive resources towards attending to the targeted speech source, in addition to processing and storing auditory information (Rönnberg et al., 2013; Pichora-Fuller et al., 2016; Peelle, 2018). This is especially concerning for individuals with hearing loss who on average report significantly increased listening effort and fatigue as scored through subjective measures and indirect measures of brain activity (McGarrigle et al., 2014). Along with the innate loss of spectral resolution due to the CI processor, the demand of comprehending degraded speech places additional strain on the cognitive resources of CI users and can lead to decreased motivation and effort in engaging in social situations (Pichora-Fuller et al., 2016; Castellanos et al., 2018). Perceived cognitive demand and listening effort has been found to be a primary factor that is detrimental to QoL amongst individuals with hearing loss, especially in CI users (Hughes et al., 2018; Holman et al., 2019).

Current understanding of neural processes underlying the management of cognitive resources when listening in complex environments is limited. Including neural speech tracking, various methods have been devised to measure cognitive demand at a neural level. Alpha oscillations (neural oscillations between 8–12 Hz) as a potential neural marker of effort and demand has been studied extensively (Foxe and Snyder, 2011; Klimesch, 2012; Obleser et al., 2012; Strauß et al., 2014; Wostmann et al., 2015; Petersen et al., 2015; Dimitrijevic et al., 2019; Magosso et al., 2019; Price et al., 2019; Hauswald et al., 2020; Hjortkjær et al., 2020; Paul et al., 2021). Increased alpha power in adverse listening conditions is hypothesized to represent the suppression of neural responses in brain regions unrelated to the task at hand (Payne and Sekuler, 2014; Strauß et al., 2014; Wilsch et al., 2015) and may reflect a conscious effort to ignore task-irrelevant stimuli (Petersen et al., 2015). Indeed, an increase in parietal alpha power has been observed during working memory tasks in individuals with hearing loss when both background noise and task difficulty increases (Petersen et al., 2015; Paul et al., 2021). A positive relationship between listening effort and alpha in the left inferior frontal gyrus (IFG) was also observed in CI users performing a digits speech-in-noise task, suggesting that language networks are involved in self-perceived listening effort (Dimitrijevic et al., 2019). When combined with visual stimuli, an increase in alpha power is seen in the parieto-occipital region when individuals attend to audio that is incongruent to the presented visual cues, suggesting a suppression of the unrelated incoming visual stimuli (O’Sullivan et al., 2019). Therefore, alpha power potentially reflects a gating mechanism towards incoming sensory stimuli in a manner relevant to task goals and may serve as a marker of cognitive demand during listening in noisy environments.

While previous literature links alpha power and speech tracking to cognitive demand, demand appears to affect speech tracking, cognitive engagement, and alpha oscillatory power differently, with speech envelope coherence demonstrating a inverted-U pattern while alpha power declines overall (Hauswald et al., 2020). Furthermore, it is unclear whether neural speech tracking and alpha power reflect self-perceived cognitive demand compared to current clinical speech perception measures. Therefore, the current objectives of the study are to first measure neural speech tracking in CI users using real-world speech, and then evaluate the effects of increasing background noise and the addition of visual cues on the quality of neural speech tracking. Then, we will quantify the relationship of neural speech tracking to the subjective self-reported level of cognitive demand and self perceived word/conversation understanding. Based on previous research regarding effort-related alpha oscillations, we conducted a secondary analysis of alpha band power over the superior-parietal and left IFG regions. We hypothesize that:

1. Self-rated cognitive demand during speech listening will decrease with the presence of visual cues but increase in response to background noise levels
2. The degree of speech tracking will be reduced by increasing background noise levels and enhanced by the presence of visual cues. Similarly, the presence of visual cues will result in a decrease in alpha power.
3. Neural speech tracking and alpha oscillatory power will be related to subjective self-reports of cognitive/mental demands and words/conversation understanding.

## 2 Materials and methods

### 2.1 Participants

Participant inclusion criteria consisted of adult CI users between the ages of 18 to 80 that have native or bilingual fluency in English, with at least 1 year of experience with their implants. As the current objective is to quantify the relationship between clinical speech perception tests and self-perceived cognitive demand in a general CI user population, potential participants were not excluded based on etiology of hearing loss, age of hearing loss onset, performance on clinical speech perception tests, or sidedness of implantation (Table 1). Eighteen adult CI users were recruited from Sunnybrook Health Sciences Centre Department of Otolaryngology. Three participants were excluded due to poor electroencephalogram (EEG) recording quality and unavailable clinical hearing scores, leaving a final participant sample of fifteen subjects (9 males, 6 females). The remaining 15 participants were between 36 to 74 years of age (M = 59.1, SD = 12.0; Table 1). The participant sample varied in terms of device implantation, and included 3 unilateral, 6 bimodal (hearing aid & CI), and 6 bilateral CI users (Table 1). Unilateral and bimodal users were on average 55.2 years old (SD = 12.4) on the date of implant activation, while the average age of bilateral CI users on the implant activation date for the ear they used for the study was 50.5 years old (SD = 13.5). All methods and protocols used in the current study were approved by the Research Ethics Board at Sunnybrook Heath Sciences Centre (REB #474-2016) in accordance with the Tri-Council Policy Statement: Ethical Conduct for Research Involving Humans. All participants provided written informed consent and received monetary compensation and full reimbursement for parking at the hospital campus for their time and participation.

**Table 1:** Demographics of cochlear implant listeners included in the current study.

| ID | Age | Sex | Implant Side | Ear Used | CI Use (yr) | Processor | Strategy | Etiology |
| --- | --- | --- | --- | --- | --- | --- | --- | --- |
| CI01 | 62 | F | Right | Right | 5 | Harmony | HiRes Optima S | Unknown |
| CI02 | 36 | M | Bilateral | Right | 6 | SONNET | FS4 | Cogan's Syndrome |
| CI03* | 53 | F | Right* | Right | 8 | OPUS 2 | FS4-P | Unknown |
| CI04* | 62 | M | Left* | Left | 2 | Naída Q90 | HiRes Optima S | Hereditary |
| CI05* | 74 | F | Right* | Right | 5 | OPUS 2 | FS4-P | Hereditary |
| CI06 | 63 | M | Bilateral | Left | 4 | Naída Q90 | HiRes Optima S | Meniere's disease |
| CI07 | 62 | M | Bilateral | Left | 2 | SONNET | FS4-P | Unknown |
| CI08 | 60 | M | Bilateral | Left | 16 | Naída Q70 | Unknown | Hereditary |
| CI09* | 44 | F | Left* | Left | 1 | SONNET | FS4-P | Unknown |
| CI10 | 71 | M | Bilateral | Right | 5 | OPUS 2 | FS4-P | Unknown |
| CI11 | 60 | F | Bilateral | Right | 11 | Freedom | ACE | Congenital |
| CI12 | 38 | F | Left | Left | 1 | SONNET | FS4-P | Meningitis |
| CI13 | 73 | M | Left | Left | 8 | RONDO | FSP | Meniere's disease |
| CI14* | 72 | M | Left* | Left | 1 | SONNET | FS4-P | Unknown |
| CI15* | 56 | M | Right* | Right | 6 | SONNET | Unknown | Unknown |

\*Participant used a hearing aid on the non-implanted side in daily life.

### 2.2 AzBio performance

Clinical speech perception was evaluated using the AzBio Sentence Test (Spahr et al., 2012). During testing, the 20-sentence lists were presented in quiet, and in noise at an SNR of +5 dB. Testing took place during clinical visits and were administered by audiologists as needed. During testing, participants were seated in a sound-isolated room with minimal reverberation, 1 meter away from a loudspeaker positioned directly in front of the participant at approximately the level of a typical listener’s head (Minimum Speech Test Battery (MSTB) For Adult Cochlear Implant Users, 2011). Participants were required to repeat back the sentence heard, and each sentence was scored as the number of words correctly reported. Percent scores are calculated for each sentence based on the percentage of words correct within the presented sentence. Sentence scores were averaged and used as the estimate of speech perception ability within the listening situation. For the current study, the most recent post-implantation test scores relative to the study date for the stimulated ear of participant (or both ears if individual ear scores were unavailable) were collected (Table 2). The average AzBio scores were 86.9 for listening in quiet (SD = 8.51), and 60.2 for listening in noise (SD = 26.6).

**Table 2:** AzBio performance of CI listeners in quiet and in noise (SNR +5dB) for the ear used during listening task. The side used during the study is listed as Listening Side. Listening scores for both ears are reported for two participants as their scores for individual ears were not available.

| ID | Listening Side | Tested Ear | Score in Quiet (%) | Score in Noise (%) |
| --- | --- | --- | --- | --- |
| CI01 | Right | Right | 88 | 22 |
| CI02 | Right | Right | 97 | 95 |
| CI03* | Right | Right | 74 | 29 |
| CI04* | Left | Left | 71 | 20 |
| CI05* | Right | Both | 92 | 50 |
| CI06 | Left | Left | 95 | 56 |
| CI07 | Left | Both | 80 | 31 |
| CI08 | Left | Left | 90 | 69 |
| CI09* | Left | Left | 79 | 60 |
| CI10 | Right | Right | 95 | 95 |
| CI11 | Right | Right | 83 | 70 |
| CI12 | Left | Left | 93 | 94 |
| CI13 | Left | Left | 79 | 49 |
| CI14* | Left | Left | 91 | 74 |
| CI15* | Right | Right | 96 | 89 |

\*Participant used a hearing aid on the non-implanted side in daily life.

### 2.3 Continuous listening paradigm

#### 2.3.1 Listening environment

The continuous listening paradigm took place in a sound-attenuated and electrically shielded booth with minimal reverberation. Participants were seated at the center of the booth, surrounded by a circular ring array of 8 speakers located at 0°, ± 45°, ± 90°, ±135°, and 180°. The speaker ring array was raised 1 meter from the ground, with each speaker cone located 0.80 meters away from the center of the ring. A computer monitor was placed 0.80 meters directly in front of the listener’s head, under the speaker located at 0° azimuth. Audio stimuli were played from the loudspeaker at 0° azimuth located directly in front of the listener, while multi-talker background noise was played from surrounding speakers.

#### 2.3.2 Audiovisual stimuli

The stimuli used for the paradigm were 15-minute audio and video segments of episodes one to four from the first season of *The Office* television show. The show was chosen as it features naturalistic dialogue with little to no music, as well as its use in previous neuroimaging studies (e.g. Byrge et al., 2015; Pantelis et al., 2015). In addition, the opening sequence containing the theme song of the show was removed to minimize non-speech sounds. To investigate cognitive demand in complex listening conditions, segments of the show were presented in conditions that varied in multi-talker babble noise levels. The babble noise soundtrack used for the current study was adapted from the four-talker babble noise used in the Quick Speech in Noise (QuickSIN) test (Killion et al., 2004).

Prior to the start of the paradigm, bilateral and bimodal CI participants were asked to remove the cochlear implant or hearing aid that was contralateral to the ear chosen for listening. Participants were asked to pay attention to the television show, attending to the audio and video stimuli presented from the speaker and monitor located directly in front of the participant at the 0° azimuth. While participants attended to the target stimuli, babble noise was simultaneously presented from the seven speakers located at the +/− 45°, +/− 90°, +/− 135°, and 180° azimuths. Each episode was presented in blocks of 5-minute segments, with each segment varying in background noise level. Movie audio segments were presented at 65 dB SPL, while the babble noise was varied to achieve the desired signal-to-noise ratio (SNR). In the Low-Noise condition, the SNR of the audio-video segment to the background babble noise was +15 dB. Audio-video segments were presented in the Moderate-Noise and High-Noise conditions at +10 dB and +5 dB SNR, respectively. Finally, to account for the benefits of visual cues for speech tracking, segments of only audio were presented in the Audio-Only condition at SNR+15 dB, with the video being replaced by a visual crosshair that served as a visual fixation point. Three runs for each condition were performed for a total of 15-minutes per condition, with the order of runs and conditions randomized for each participant to avoid bias. The presentation of video and audio segments were processed in MATLAB 2009b (The MathWorks, Natick, MA, USA) and controlled by a Tucker Davis Technologies (TDT, Alachua, FL, USA) RX8 Processor. After each episode, participants were asked to answer nine simple open-response questions on the events that occurred in the episode to ensure that they attended to the stimuli, averaging to 79.63% correct (SD = 19.53%).

#### 2.3.3 Assessment of self-perceived listening effort

In order to measure self-reported listening effort, participants responded to the first section of the NASA-Task Load Index (Hart and Staveland, 1988) after each 5-minute run. The NASA-Task Load Index (NASA-TLX) is a multi-scale self-report that rates an individual’s perceived workload for a task (Hart and Staveland, 1988). The dimensions consist of six ordinal subscales: Mental Demand, Physical Demand, Temporal Demand, Performance, Effort, and Frustration. For the current study, the scales for Physical Demand and Temporal demand were removed. The Effort subscale was merged with Mental Demand in order to align with the concept of listening effort as defined as the mental energy an individual perceives they need to meet external demands (Pichora-Fuller et al., 2016; Peelle, 2018). Specifically, participants responded to the question “How mentally demanding was the task?” on a 21-point scale to gauge cognitive demand, with the endpoints on the left and right being “Low” and “Very High”, respectively. Additionally, subscales assessing the perceived percentage of words understood and the percentage of conversation understood were added.

### 2.4 Electrophysiological recording

EEG data were recorded using a 64-channel antiCHamp Brain Products recording system (Brain Products GmbH, Inc., Munich, Germany) at a sampling rate of 2000 Hz. EEG caps were fitted onto the participants so that the electrode corresponding to the typical Cz location according to the 10-20 international system was at the vertex of the skull, as determined using the point of intersection between nasion-to-inion and tragus-to-tragus midpoints. The EEG online reference was a separate reference electrode located slightly anterior to the vertex on the midline, while the ground electrode was located on the midline at the midpoint between the nasion and the vertex. Electrodes in proximity to or overlapping the CI magnet and coil on the side of the stimulated ear were not used; this ranged from 1 to 3 electrodes across all participants.

#### 2.4.1 EEG preprocessing

EEG recordings were imported into EEGLAB (Delorme and Makeig, 2004) for MATLAB 2019a (The MathWorks, Natick, MA, USA) before being filtered with a Finite Impulse Response bandpass filter from 0.3 to 40 Hz. Stimuli audio segments were simultaneously recorded along with the EEG in an auxiliary channel to maximize the temporal alignment of the audio stimuli to the EEG data. The audio soundtrack was then aligned offline to the recorded audio stream by calculating the cross-covariance to find the corresponding starting time point of the soundtrack itself. Speech envelopes were extracted by calculating the absolute value of the Hilbert transformation of the aligned audio soundtrack. The extracted envelope was then downsampled to the EEG recording sampling rate and appended to the EEG data for later analyses. After alignment, the EEG and audio data were both downsampled to 250 Hz and concatenated together across runs and conditions for each participant. Noisy channels and durations of extremely noisy data were removed prior to artifact removal. Independent Component Analysis (ICA) was conducted on the concatenated data to identify stereotypical physiological artifacts (e.g., eye blinks, oculomotor movements, cardiac activity) and technical artifacts (e.g., electrode pop, line noise). Components containing these artifacts were manually removed based on the visual inspection of component topographies. An average of 6.53 (SD = 1.13) components were removed across all participants. The recordings were then separated based on study condition for each participant, and the previously removed channels were interpolated using the spherical splines of neighbouring channels (Perrin et al., 1989).

#### 2.4.2 CI artifact suppression

In the presence of auditory stimuli, the electrical stimulation and radio-frequency signals of CIs impart electrical stimulation artifacts into the EEG recording (Wagner et al., 2018). While ICA has been previously used to identify CI artifacts in EEG recordings involving auditory evoked potentials (Gilley et al., 2006; Miller and Zhang, 2014), the nature of CI artifacts makes their reduction more challenging, especially for continuous tasks. In general, ICA separates statistically independent components from the mixed signal. In the case of artifact reduction in MEG and EEG data, temporal ICA separates signal components based on their temporal independence. Artifacts that are not temporally aligned to the stimulus onset (e.g., eye blinks), are therefore separated from sources that are temporally aligned (e.g., event-related potentials). CI artifacts however, are temporally aligned to the sound onset in a continuous EEG recording, and have a similar morphology to the sound envelope (Mc Laughlin et al., 2013).

Second-order blind identification (SOBI) was applied (Belouchrani et al., 1997) to reduce CI-related artifacts as we have used previously (Paul et al., 2020; Paul et al., 2021). As the name suggests, SOBI is a blind source separation technique that uses second-order statistics in order to separate temporally correlated sources (Belouchrani et al., 1997). SOBI was performed on the continuous EEG data for each participant, and components of suspected CI artifacts were visually identified based on the topographical centroid and implant side used during the listening paradigm. Between 0 and 2 components corresponding to the artifact centroids were removed for each participant (M = 1.58, SD = 0.62). After CI artifact suppression, the cleaned EEG and aligned audio stimuli were subsequently imported into Fieldtrip (Oostenveld et al., 2011) and re-referenced to an average reference before further processing to prepare for TRF and alpha power analysis.

### 2.5 Data analysis

#### 2.5.1 TRF calculation

TRFs were estimated from EEG data and the speech envelope of the audio signal using the mTRF Toolbox 2.0 (Crosse et al., 2016) in MATLAB. The processed audio signal and the EEG signal were filtered by a 1 to 20 Hz bandpass, 2^nd^ order, zero-phase filter. The filtered EEG and audio signals were subsequently z-scored for each participant before TRF calculations. Integration window of time lags between −100 to 500 ms were chosen for analysis. The *mTRFcrossval* function was used to optimize the regularization parameter that reduces the overfitting of the speech envelope to the EEG data. The *mTRFcrossval* function performs a leave-one out *N*-fold cross-validation, where for each participant TRFs are estimated for all trials except one. The estimated encoding model is then used to predict the EEG signal for the trial that was omitted, and the Pearson’s correlation coefficient is calculated to assess the correlation between the predicted and actual EEG signals. This process was repeated for regularization parameters ranging from 2^-5^ to 2^15^.

While some studies suggest the use of uniquely optimized regularization parameters for each participant dataset to maximize the TRF estimation (Crosse et al., 2021), this can potentially overfit the TRFs to remnant CI artifacts in CI user recordings. Thus, the correlation coefficients were then averaged across all subjects, channels, and conditions for each regularization parameter, and the value associated with the highest correlation value was chosen as the optimal parameter to use. The continuous EEG data was therefore epoched into 60-second trials prior to TRF calculation, and 2^11^ was selected as the regularization parameter. TRF epochs were averaged within each condition for every participant, generating four sets of TRFs per participant for each listening condition.

#### 2.5.2 Neural speech tracking analysis

Prior to analysis, TRFs for each channel were baseline corrected by subtracting the mean of the values between the baseline period (−100 to −8 ms) from each point in the TRF waveform using Brainstorm (Tadel et al., 2011). Previous literature has identified that speech tracking component amplitudes reaching their maximum magnitudes within the frontocentral region (e.g. Ding & Simon, 2012; Fiedler et al., 2019). Thus, we focused on the frontocentral scalp region as the region of interest (ROI) at the sensor level. Specifically, the TRFs for seven frontocentral EEG sensors within the chosen ROI were averaged for each participant per condition to generate a mean TRF for further analysis (Figure 1A).

**Figure 1:**
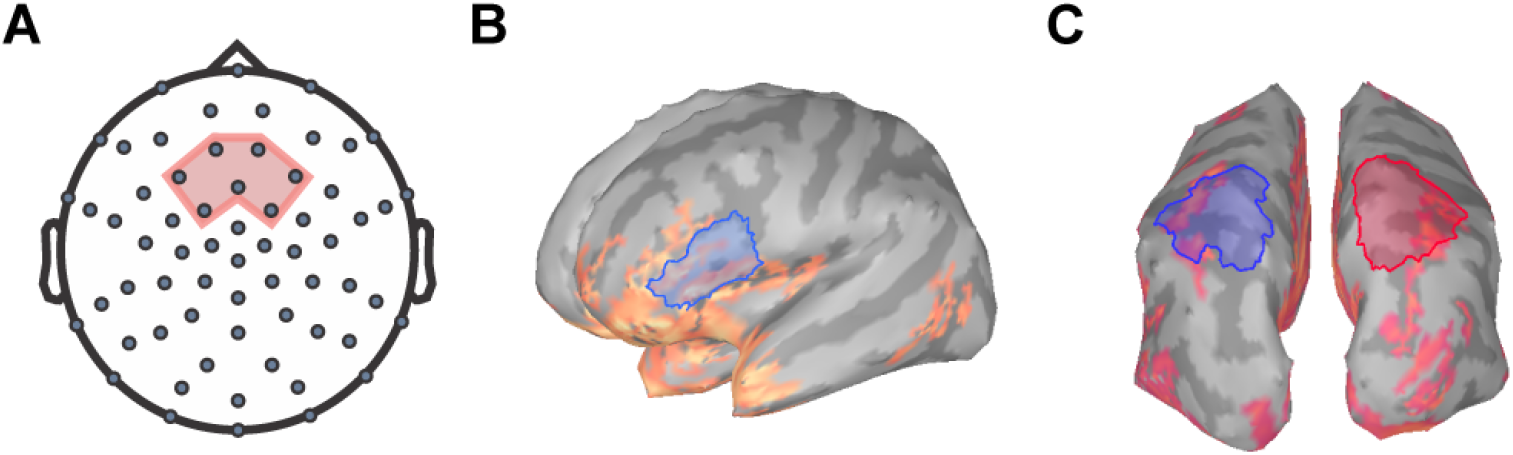
Regions of interest for TRF and frequency analysis. The 7 frontocentral channels EEG channels (**A**) chosen for TRF analysis are within the region shaded in red. Relative alpha power was extracted at the (**B**) left inferior-frontal gyrus and the (**C**) superior-parietal regions for the both hemispheres (left in blue, right in red).

To identify the time windows of interest containing the relevant speech components, the babble noise audio was used to create a comparison standard with similar spectral but a temporally different speech envelope. As with the stimuli soundtrack, the absolute value of the Hilbert transformation was calculated for the babble comparison soundtrack to extract the amplitude envelope. The comparison soundtrack was then scaled to the root-mean-squared amplitude of attended soundtrack amplitude before being subjected to the same TRF estimation procedure. TRF differences between the stimuli TRF and the babble noise TRF at each time lag were first compared using two-tailed paired t-tests. T-test values were then corrected using the Benjamini-Hochberg procedure (Benjamini and Hochberg, 1995) was used to correct for repeat testing. TRF differences at time lags were considered significantly different if the adjusted p-values were less than the alpha criterion of 0.05. Component TRF peak amplitudes and peak latencies were then extracted by finding the point of the local maxima/minima of the component within the identified time lag windows.

#### 2.5.3 Alpha oscillatory power

The involvement of the IFG in neuronal speech tracking has been examined in various studies (e.g., Zion Golumbic et al., 2013b; Price et al., 2019). We have previously shown that neural tracking within the left IFG has been observed to be correlated with listening effort and predicted successful trials for speech perception (Dimitrijevic et al., 2019). Additionally, previous literature has indicated that parietal alpha oscillations may reflect cross-modal attentional modulation (e.g. Misselhorn et al., 2019). Relative alpha power was extracted from EEG data using Brainstorm following the methods outlined in Niso et al., 2019. The preprocessed spontaneous EEG recordings were first imported into Brainstorm, and source activations were calculated using standardized low-resolution electromagnetic tomography (sLORETA) modeling (Pascual-Marqui, 2002). sLORETA models provide estimates of the power and location of the neural generators that underlie electrophysiological processes. Boundary element model (BEM) head models were created using the OpenMEEG plugin prior to sLORETA modeling. sLORETA models were z-scored, and alpha power was computed using the Brainstorm resting-state pipeline as outlined in Niso et al., 2019.

The power spectral densities (PSDs) of each voxel of the sLORETA models were estimated from 0 to 125 Hz by applying Welch’s method using a window of 1 second at 50% overlap to obtain a 1.0 Hz frequency resolution. Spectrum normalization was then applied by dividing the PSDs by the total power. PSDs were then averaged across all participants and sides for each condition to identify ROIs for alpha analysis. Alpha power was extracted over the left IFG ROI, a 38.74 cm^2^ region encompassing the pars triangularis and the pars opercularis (Figure 1B) as defined by the Desikan-Killiany Atlas (Desikan et al., 2006). Additionally, ROIs encompassing the superior-parietal cortices (Desikan et al., 2006) were also defined as a 57.32 cm^2^ area within the left posterior parietal region and a 57.45 cm^2^ area within the right posterior parietal region (Figure 1C). Mean relative alpha power was calculated by averaging alpha power values between 8 to 12 Hz and then dividing by the total power. The relative alpha power was then extracted over all ROIs for each participant per condition.

### 2.6 Statistical analysis

All statistical analyses were performed in R (R Core Team, 2021) and MATLAB 2019a (MATLAB, 2019). One-way repeated measures analysis of variance (RM-ANOVA) with the main effect of Condition (Low-Noise, Moderate-Noise, High-Noise, Audio-Only) were used to compare behavioural scores. Two-way factorial RM-ANOVAs with the main effect of Condition and TRF Component (TRF_100_, TRF_200_, TRF_350_) were performed to compare the effect of background noise and visual cues on TRF component amplitudes. Two-way RM-ANOVAs with the main effects of Condition and Listening Side (Ipsilateral to implant, Contralateral to implant) and a paired t-test were performed to compare relative alpha power for the superior-parietal and IFG ROIs, respectively. The relationship between listening demand and neural variables were analyzed separately for TRF components and relative alpha power. In cases where sphericity is violated, Greenhouse-Geisser corrections were applied to p-values (p_GG_). For all tests, the alpha criterion was set at 0.05, and p-values (p_adj_) were adjusted for multiple comparisons using the Benjamini-Hochberg procedure (Benjamini and Hochberg, 1995). Generalized eta squared (η^2^_G_) values are also reported alongside ANOVA test statistics to indicate effect size.

Linear mixed models were applied to predict listening demand, perceived percentage of words understood, and perceived percentage of the conversation understood from the fixed effects of TRF component or relative alpha power, listening condition, and subject age using the R package *lme4()* v1.1.29 (Bates et al., 2015). TRF component amplitudes, alpha power values, and age were treated as continuous variables and z-scored prior to modeling. Although by-subject random effects and condition were included as random intercepts and random slopes respectively to account for the variability explained by the aforementioned factors, random effects are not interpreted due to being jointly unidentifiable as each subject was statistically measured once per condition. Additionally, the by-subject random slope for condition was removed from the perceived percentage of words models due to nonconvergence. The correlation between the random intercept for subject and the by-subject random slope for condition was removed for the perceived percentage of conversation models due to convergence issues as well. Fixed effects were subsequently analyzed using ANOVAs with degrees of freedom adjusted by the Satterthwaite method (Luke, 2017), and all post-hoc analyses were performed through pairwise comparisons of the estimated marginal means. As there were no AzBio score matching the SNR+15 dB and +10 dB conditions, two-tailed partial Pearson correlations while controlling for age were used to correlate neural measures and AzBio scores. The relationship between perceived percentage of conversation understood and AzBio scores were also explored using a two-tailed partial Spearman correlation while controlling for age. For the partial correlations, the alpha criterion was set at 0.05 and p-values (p_adj_) were adjusted for multiple comparisons using the Benjamini-Hochberg procedure (Benjamini and Hochberg, 1995).

## 3 Results

### 3.1 Self-reported listening demand, percentage of words and conversation understood

Average self-reported NASA-TLX mental demand ratings during listening in the audiovisual conditions were 6.78 (SD = 2.58) for the Low-Noise (LN) condition, 11.21 (SD = 3.20) for the Moderate-Noise (MN) condition, and 15.44 (SD = 2.61) for the High-Noise (HN) condition. The average mental demand rating for the Audio-Only (AO) condition was 14.62 (SD = 2.63). The ANOVA model returned an effect of condition on mental demand ratings (F_(3,42)_ = 40.83, p < 0.001, ƞ^2^G = 0.620).

As expected, self-reported mental demand ratings increased as background babble noise levels increased during the audiovisual listening paradigm (Figure 2A). Post-hoc analysis using paired t-tests revealed that the average demand rating was significantly higher in the HN condition compared to the MN condition (p_ad j_< 0.001), and the LN condition (p_adj_ < 0.001). Additionally, the MN condition demand rating was also statistically significantly higher than for LN (p_adj_ = 0.007). To compare the effects of visual cues on listening effort, the background babble noise level for the LN and AO conditions were both set at SNR+15 dB. Post-hoc analysis indicates that the average demand rating for AO was significantly higher than LN (p_adj_ < 0.001) despite taking place in the same background noise level. Interestingly, while the AO demand rating was statistically significantly higher compared to the MN rating (p_adj_ = 0.009) as well, it was not significantly different compared to the HN demand rating (p_adj_ = 0.322) despite the 10 dB difference in SNR between the two listening conditions.

**Figure 2:**
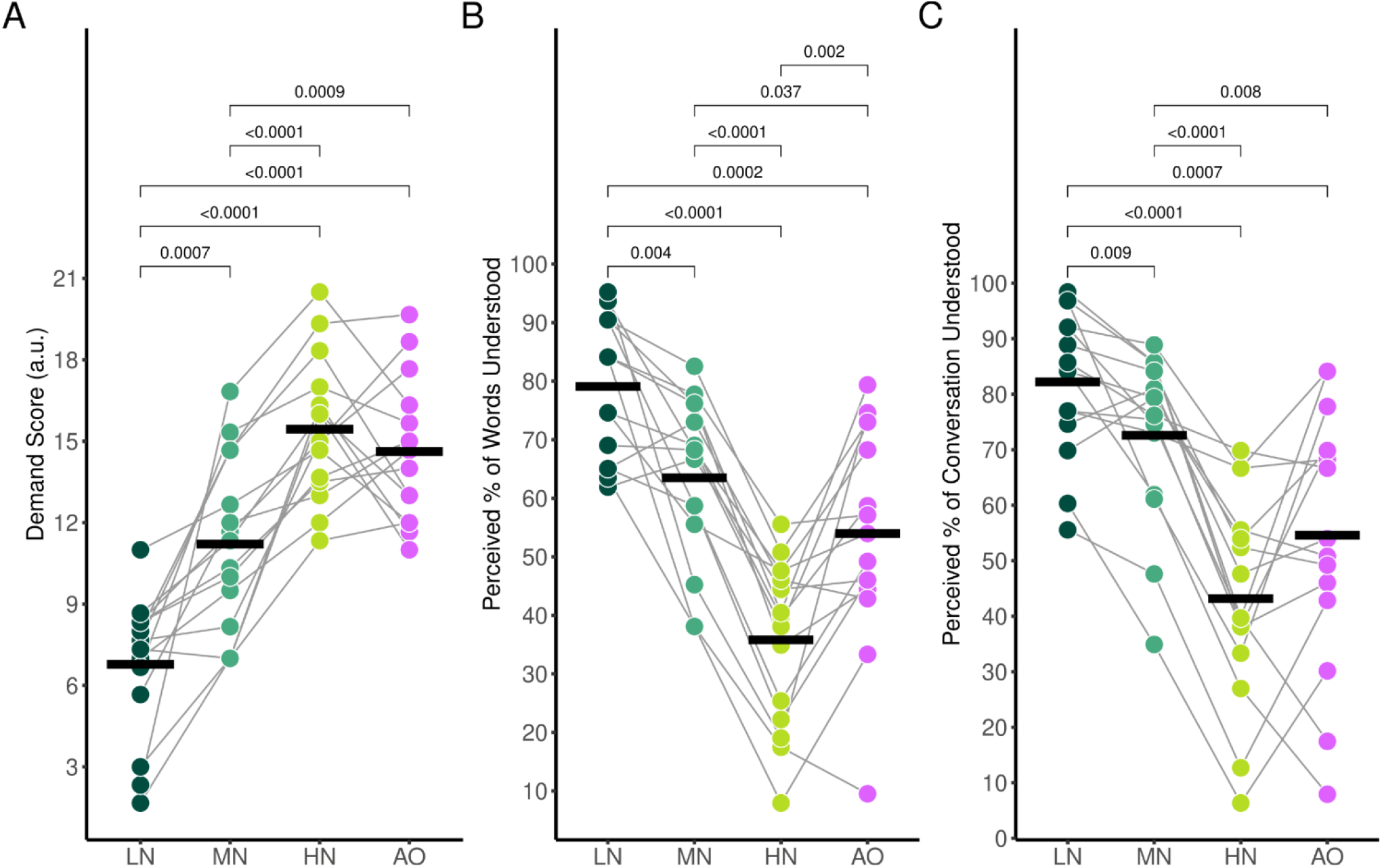
Average self-reported NASA-TLX ratings for (**A**) mental demand, (**B**) perceived percentage of words understood, and (**C**) perceived percentage of the conversation understood. Black horizontal bars indicate the mean score of each condition. Only significant comparisons are displayed.

The average percentage of words understood followed the same trends (Figure 2B); participants reported 79.1% (SD = 12.0%) of words understood for the LN condition, 63.5% (SD = 14.1%) for the MN condition, 35.8% (SD = 14.1%) for the HN condition, and 54.0% (SD = 18.5%) for the AO condition. There was a significant effect of condition (F_(3,42)_ = 36.67, p_GG_ < 0.001, η^2^_G_ = 0.543), with all conditions significantly differing as shown by post-hoc pairwise analysis. The percentage of words understood in the LN condition was significantly higher compared to the MN (p_adj_ = 0.004). HN (p_adj_ < 0.001), and AO (p_adj_ < 0.001) condition. The MN percentage rating was significantly higher than the HN (p_adj_ < 0.001) and AO (p_adj_ = 0.037) ratings as well. The percentage of words understood in the HN condition was significantly lower compared to the percentage understood in the AO condition (p_adj_ = 0.002).

Perceived percentage of conversation understood also decreased parametrically as background noise increased (Figure 2C). Participants reported that they understood an estimated 82.2% (SD = 13.1%) of the conversation for the LN condition, 72.6% (SD = 15.2%) for the MN condition, 43.2% (SD = 18.7%) for the HN condition, and 54.6% (SD = 23.1%) for the AO condition. The ANOVA model revealed a significant effect of condition (F_(2.10,29.36)_ = 27.65, p_GG_ < 0.001, η^2^_G_ = 0.435). Post-hoc analysis revealed that the perceived percentage of the conversation understood differed across all conditions. The percentage rating in the LN condition was significantly higher compared to the MN (p_adj_ = 0.009), HN (p_adj_ < 0.001), and AO conditions (p_adj_ = 0.007). The rating in the MN condition was also higher than the HN (p_adj_ < 0.001) and AO condition (p_adj_ = 0.079). Similar to mental demand ratings, the percentage of words understood also did not significantly differ between the HN and AO conditions (p_adj_ = 0.051).

### 3.2 Effect of increasing background noise

#### 3.2.1 Effect on TRF components

Movie audio-driven TRFs at frontocentral sensors in conditions of increasing background noise resembled N1-P2 cortical auditory evoked potentials (Figure 3A) has been observed in past studies (Hjortkjær et al., 2020; Verschueren et al., 2020). Comparison of TRFs to the movie audio and babble noise in Figure 3B revealed two significant time lag windows containing negative TRF components for the LN and MN conditions, where movie-audio TRFs differed from babble noise TRFs ~100 ms time lag and ~ 350 ms time lag, herein referred to as TRF_100_ and TRF_350_ respectively. Visual inspection of the TRF at frontocentral sensors also suggested a TRF component at 200 ms time lag (TRF_200_) but this did not significantly differ from the TRF estimated from the background babble noise in each condition. Nonetheless the TRF_200_ was submitted to statistical analysis to compare across latencies. Based on these responses, an 8 ms time window surrounding the grand mean component peaks were chosen for analysis for the TRF_100_ and TRF_200_. A 48 ms time window surrounding the TRF_350_ component peaks was chosen due to the previously observed wider time lag window width. The average component peak amplitudes were subsequently calculated by averaging the mean stimuli TRF values within the chosen time lag intervals for each participant and compared between conditions using a two-way RM-ANOVA with the main effect of Condition (Low-Noise, Moderate-Noise, High-Noise) and TRF Component (TRF_100_, TRF_200_, TR_F350_).

**Figure 3:**
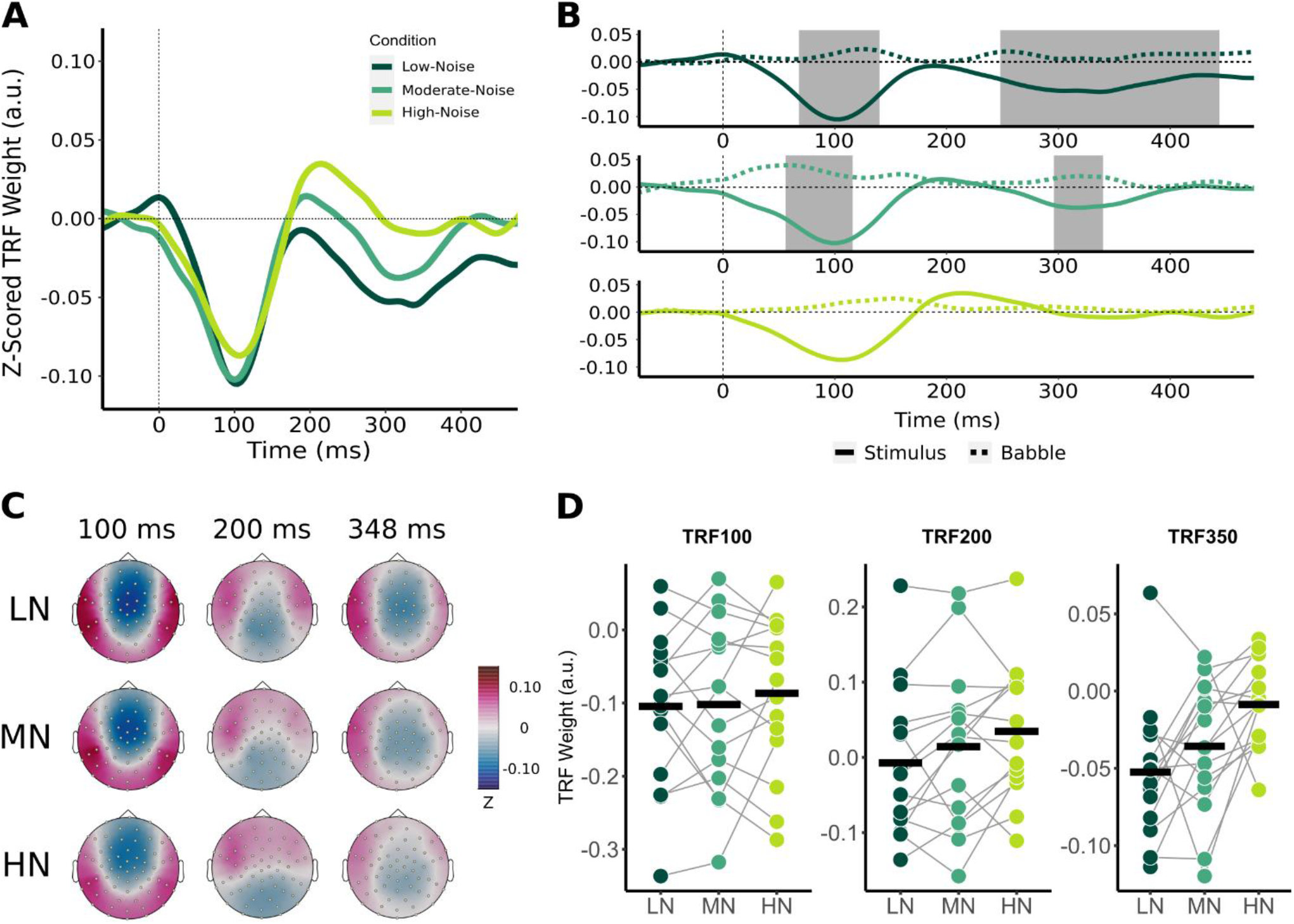
Effect of increasing background noise on sensor-level TRF peaks. (**A**) Mean condition TRFs compared between the audiovisual Low-Noise (LN), Moderate-Noise (MN), and High-Noise (HN) conditions. (**B**) Mean stimuli (solid-line) and background noise (dotted-line) TRFs are compared for each condition (LN - top; MN - middle; HN - bottom) using paired t-tests. Shaded regions indicate time intervals where the TRFs were significantly different after correcting for multiple comparisons. (**C**) Topographies of the mean condition stimuli TRFs at the 100 ms, 200 ms, and 350 ms time lags. (**D**) TRF peaks were compared between the LN, MN, and HN conditions. Black horizontal bars indicate the mean TRF weight of each condition. No statistically significant differences between TRF amplitude averages were detected.

Initially, mean TRF waveforms of the audiovisual conditions appear to differ in average amplitude for all three speech tracking components (Figure 3A, D). TRF_100_ amplitude seemed to decrease concurrently with SNR, with mean amplitudes being −0.104 (SD = 0.106) for LN, −0.102 (SD = 0.116) for MN, and −0.087 (SD = 0.107) for the High-Noise condition. While the mean TRF_200_ component was not significantly different from the babble TRF, TRF_200_ amplitude initially appeared to increase as SNR decreased. The TRF_200_ peak amplitudes were as follows: −0.007 (SD = 0.099) for LN, 0.014 (SD = 0.108) for MN, and 0.035 (SD = 0.090) for HN. TRF_350_ peak amplitudes followed a similar trend to the TRF_100_ component, in that amplitudes decreased as listening condition SNR decreased: TRF_350_ peak amplitudes were −0.052 (SD = 0.046) for LN, −0.058 (SD = 0.051) for MN, and −0.040 (SD = 0.032) for HN. While the ANOVA model revealed no significant interaction effect between condition and TRF component (F_(2.91,40.75)_ = 0.468, p_GG_ = 0.701, η^2^_G_ = 0.003), the main effect of Condition was statistically significant (F(1.78,24.95) = 3.58, p_GG_ = 0.048, η^2^ = 0.027). Post-hoc analysis however revealed no significant differences in general TRF component amplitude between conditions (all p_adj_ > 0.05).

#### 3.2.2 Effect on relative alpha power

The power spectral densities for each participant were estimated from the sLORETA models of the spontaneous EEG recordings. Subsequently, the relative alpha oscillatory power over superior-parietal cortices (Figure 4A) and the left IFG (Figure 4B) were compared between conditions. Relative alpha power at the superior-parietal cortices was analyzed with a two-way RM-ANOVA with the main effects of Condition (Low-Noise, Moderate-Noise, High-Noise) and Listening Side (Ipsilateral, Contralateral), while one-way RM-ANOVA with the main effect of Condition (Low-Noise, Moderate-Noise, High-Noise) was performed for the left IFG. Initially, there appeared to be a parametric increase in relative alpha power at the left IFG as listening condition demand increased. Subsequent analysis however revealed no statistically significant interaction effect between Condition and Listening Side for relative alpha power for all ROIs. Furthermore, the main effect of Condition and Listening Side were also not statistically significant.

**Figure 4:**
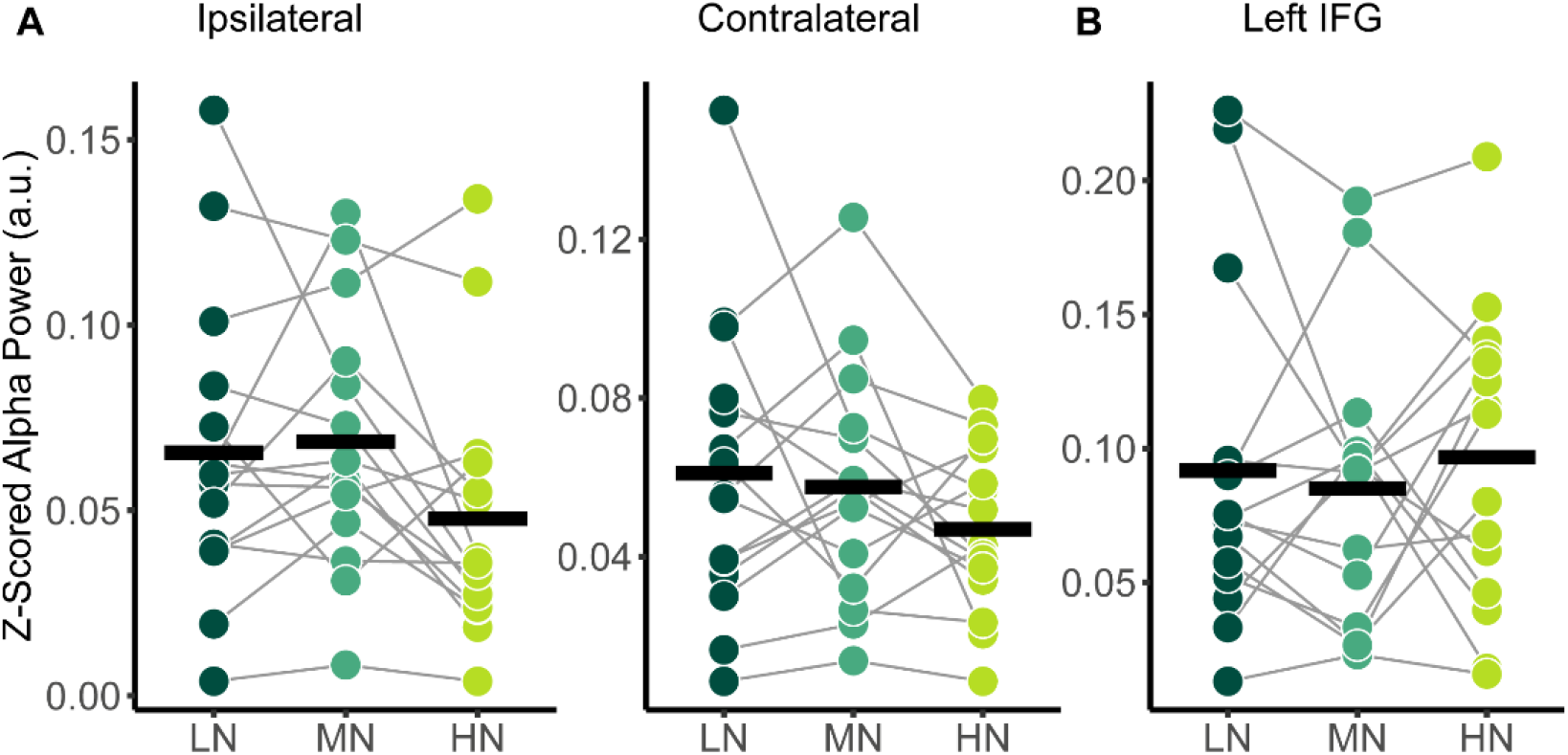
Comparison of relative alpha power between audiovisual Low-Noise (LN), Moderate-Noise (MN), and High-Noise (HN) conditions for the (**A**) ipsilateral and contralateral superior-parietal region and (**B**) left inferior-frontal gyrus (IFG) ROIs. No statistically significant differences were detected.

### 3.3 Effects of the presence of visual cues

#### 3.3.1 Effect on TRF components

Like movie audio-driven TRFs, the audio-only TRF at the frontocentral censors also resemble an N1-P2 cortical auditory evoked potential (Figure 5A) with two significant time lag windows around the 100 and 250 ms time lag (Figure 5B). As with the movie TRFs, an 8 ms time window was chosen for analysis for the TRF_100_ and TRF_200_ and a 48 ms time window was chosen for TRF_350_.

**Figure 5:**
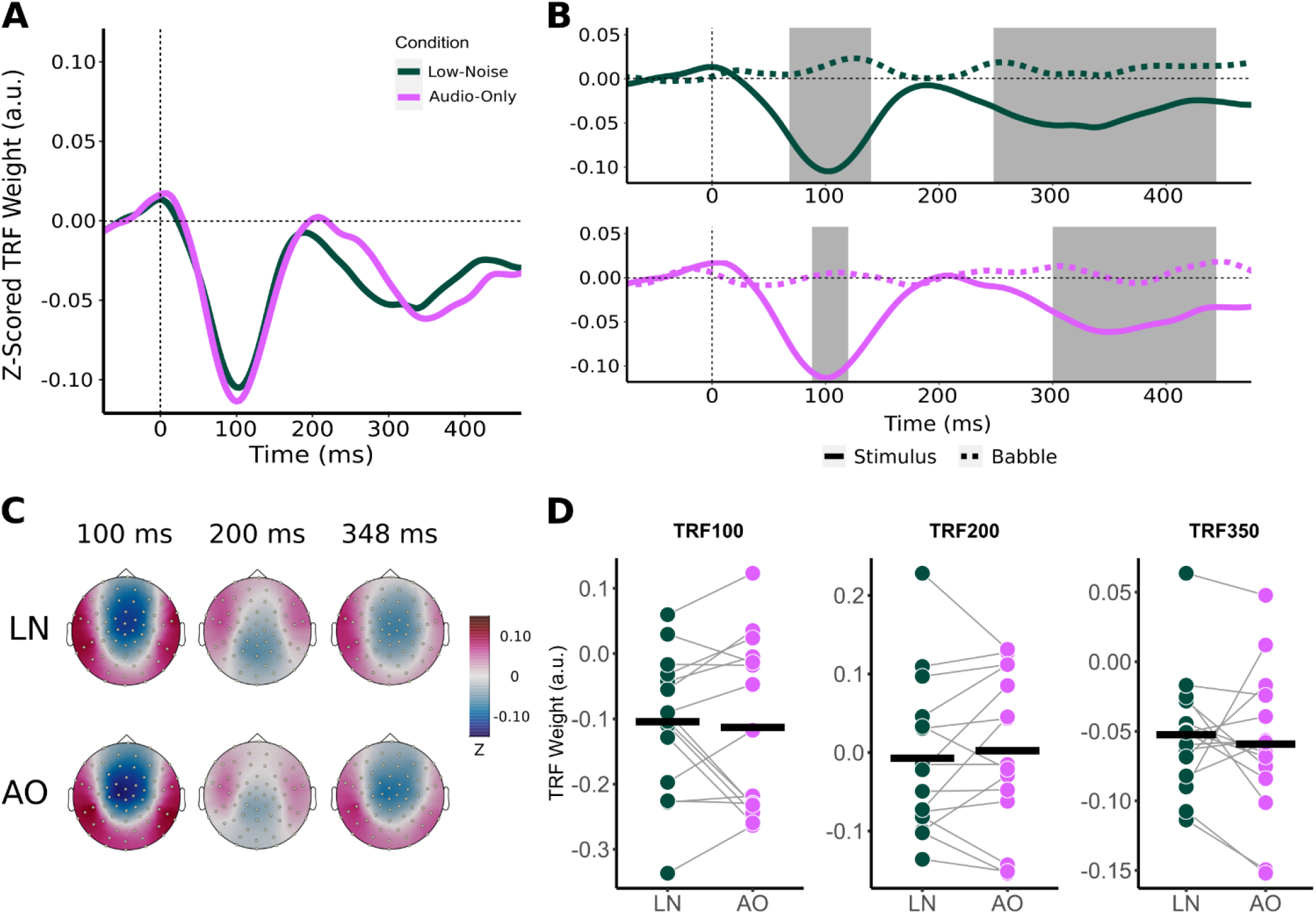
Effect of visual cues on sensor-level TRF peaks. (**A**) Mean condition TRFs compared between the audiovisual Low-Noise (LN) and Audio-Only (AO) conditions. (**B**) Mean stimuli (solid-line) and background noise (dotted-line) TRFs are compared for each condition (LN - top; AO - bottom) using paired t-tests. Shaded regions indicate time intervals where the TRFs were significantly different after correcting for multiple comparisons. (**C**) Topographies of the mean condition stimuli TRFs at the 100 ms, 200 ms, and 350 ms time lags. (**D**) TRF peaks were compared between the LN and AO conditions. Black horizontal bars indicate the mean TRF weight of each condition. No significant differences were observed.

Average component peak amplitudes were compared between LN and AO conditions using paired t-tests. Like the audiovisual conditions, TRF component amplitudes appeared to change parametrically in the more demanding AO condition (Figure 5D). Compared to LN, which had component amplitudes of −0.104 (SD = 0.106) for TRF_100_, −0.007 (SD = 0.099) for TRF_200_, and −0.052 (SD = 0.046) for TRF_350_, AO TRF components amplitudes were slightly greater. The AO TRF_100_, TRF_200_, and TRF_350_ amplitudes were −0.113 (SD = 0.132), 0.002 (SD = 0.102), and −0.059 (SD = 0.053) respectively. Similarly, no significant interaction effect between condition and TRF component was observed (F_(2,28)_ = 0.573, p = 0.570, η^2^_G_ = 0.002). The main effect of Condition was also not significant (F_(1,14)_ = 0.022, p = 0.885, η^2^ = 0.0001).

#### 3.3.2 Effect on relative alpha power

Relative alpha power was analyzed with a two-way RM-ANOVA with the main effects of Condition (Low-Noise, Audio-Only) and Listening Side (Ipsilateral, Contralateral) for the superior-parietal cortices. A paired t-test was performed for the left IFG. As with the audiovisual condition comparisons, analyses revealed no statistically significant interaction effect between Condition and Listening Side for relative alpha power the superior-parietal ROI (Figure 6A) and no significant effect of Condition at the left IFG (Figure 6B), despite the apparent parametric change in alpha with condition demand. Furthermore, the main effects of Condition and Listening Side were also not statistically significant for the superior-parietal region.

**Figure 6:**
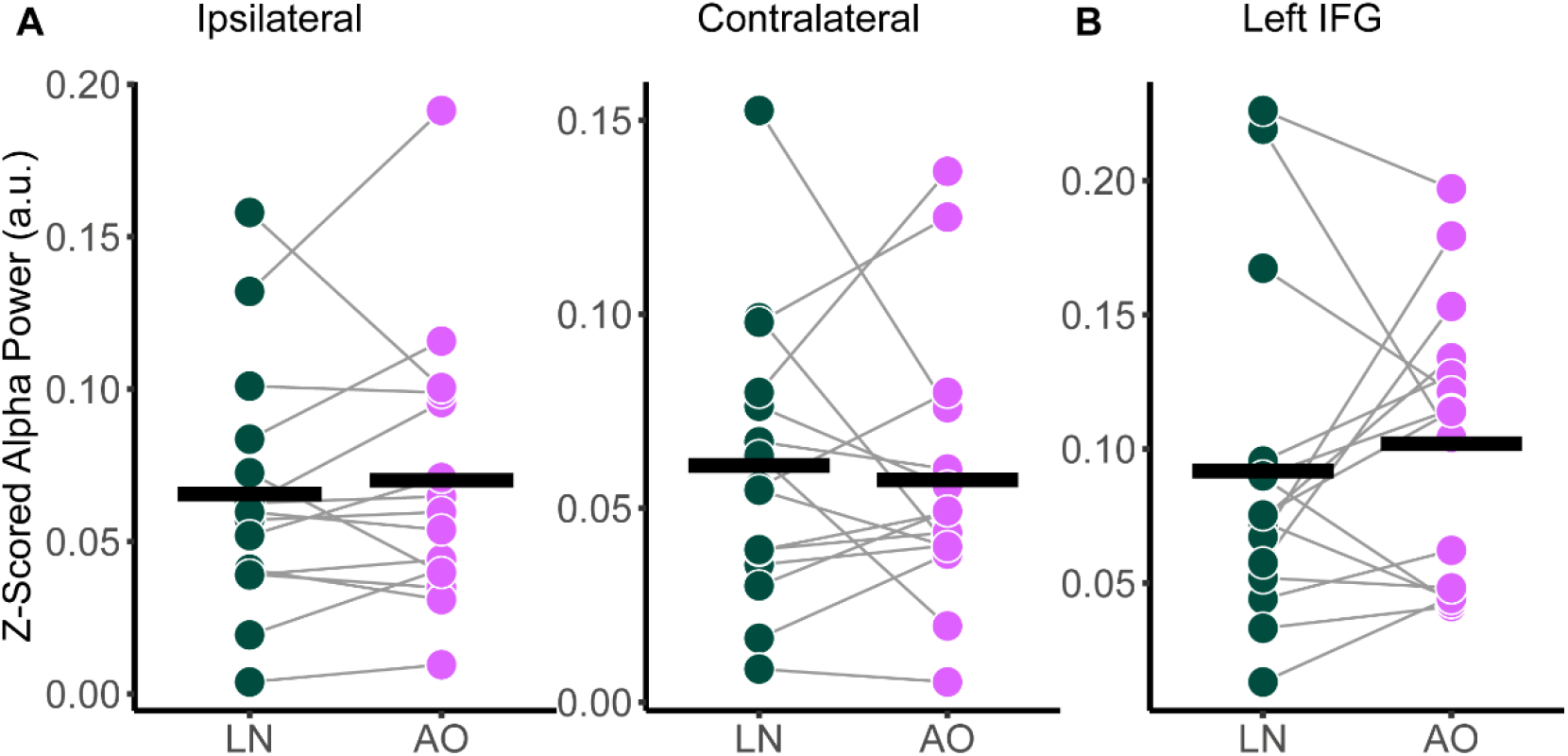
Comparison of relative alpha power between audiovisual Low-Noise (LN) and the Audio-Only (AO) conditions (bottom row) for the (**A**) ipsilateral and contralateral superior-parietal region and (**B**) left inferior-frontal gyrus (IFG) ROIs. No statistically significant differences were detected.

### 3.4 Relationship between neural measures, subjective listening demand, perceived percentage of conversation understood, and AzBio scores

Across all models, the Audio-Only condition was set as the reference category, and the fixed effect of condition was significant (all p < 0.001), supporting the previous analysis on differences in subjective behavioural scores between listening conditions. Thus, the fixed effect of condition was not analyzed further owning to the previous analyses performed.

Mixed effects modeling on self-reported mental demand during listening and TRF component amplitude while accounting for participant age indicates that demand scores increase as negative-going TRF component magnitudes attenuate. ANOVA on model fixed effects report significant effects of TRF_100_ amplitude (standardized β = 1.961, SE = 0.419, F_(1,31.4)_ = 5.461, p < 0.001) and age (standardized β = 0.646, SE = 0.267, F_(1,7.7719)_ = 5.847, p = 0.043). No other fixed effects aside from condition reached significance (all p > 0.110). Regarding the percentage of words understood, no significant fixed effects other than condition was detected (all p > 0.069). In contrast, ANOVA on the fixed effects of the mixed model on perceived percentage of conversations understood and TRF component amplitudes while accounting for age also indicates a significant main effect of TRF_100_ amplitude (standardized β = −1.418, SE = 1.058, F_(1,19.797)_ = 11.361, p < 0.001) as well as an interaction effect between condition and TRF_200_ amplitude (F_(3,13.071)_ = 3.428, p = 0.049). However, there was no specific interaction between fixed effects (all p > 0.090).

Similar models were fitted for relative alpha power at the superior-parietal and left IFG ROIs. Since there was no significant main effect of Listening Side, the superior-parietal relative alpha power at the ipsilateral and contralateral hemispheres were first averaged for each participant to create a composite measure. ANOVA on the fixed effects revealed a significant interaction between superior-parietal relative alpha power and condition (F_(3,13.364)_ = 3.457, p = 0.047), specifically in the High-Noise condition (standardized β = −1.992, SE = 0.701, p = 0.014). As with the TRF model, the fixed effect of age was also significant (standardized β = 0.6460, SE = 0.267, F_(1,7.7719)_ = 5.846, p = 0.043). For the left IFG, ANOVA on the model fixed effects reported no significant terms (all p > 0.056). The regression lines and the 95% confidence intervals of z-scored TRF_100_ amplitude against the respective self-report scores are plotted in Figure 7A and Figure 7B. Weaker TRF_100_ components displaying increasingly positive amplitudes (as TRF_100_ is a negative component) are correlated with higher demand scores (Figure 7A). In tandem, weaker TRF_100_ components are also correlated with a lower perceived percentage of the conversation understood (Figure 7B). Figure 7C shows the mean superior-parietal alpha power is plotted against demand for the High-Noise condition along with the regression line and the 95% confidence interval to show the association between increasing alpha power in the superior-parietal region and decreasing demand ratings.

**Figure 7:**
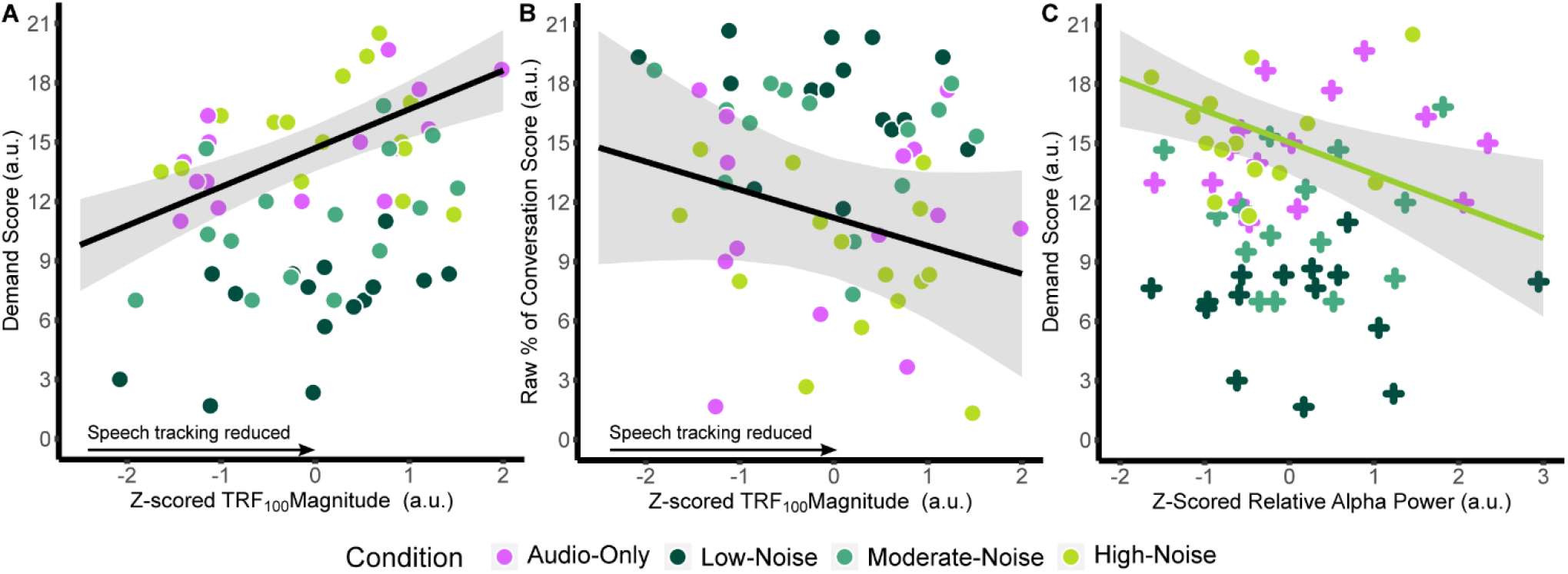
Plots of the function between (**A**) z-scored TRF_100_ amplitude and listening demand ratings, (**B**) z-scored TRF_100_ amplitude and the perceived percentage of conversation understood. The black line indicates the regression line, bounded by the 95% confidence intervals indicated in the shaded grey region. (**C**) plots z-scored relative alpha power and listening demand ratings in the High-Noise condition. The green line indicates the regression line of the High-Noise condition, bounded by the 95% confidence intervals indicated in the shaded grey region. The plus symbols indicate relative alpha power in other listening conditions.

Two-tailed partial Pearson correlations between TRF component amplitudes and AzBio In-Quiet and In-Noise scores revealed no significant correlations (all p_adj_ > 0.160). Similarly, no significant correlations were found between relative alpha power and AzBio scores across all ROIs (all p_adj_ > 0.246). Correlations between AzBio In-Quiet and In-Noise scores and perceived percentage of conversation understood also revealed that while the percentage rating correlated with AzBio In-Quiet and In-Noise scores in the AO condition prior to correcting for multiple comparisons, no correlations survived corrections for multiple comparisons (all p_adj_ > 0.075).

## 4 Discussion

The current study investigated neural speech tracking and alpha power and the degree to which these neural measures are reflected in the self-reported mental demand of listening in CI users when attending to naturalistic stimuli. We found: (1) All subjects reported higher demand ratings and lower word/conversation understanding as the noise levels increased. When participants had no video and only listened to the movie the reported demand and conversation understanding were comparable to the high noise condition even though the babble noise levels were the same as the low noise condition. (2) Neural tracking/alpha differences: The neural tracking data showed no significant parametric change in TRF components with increasing babble noise. No differences in neural tracking were observed when the movie video was replaced with a cross hair. Alpha power did not show parametric changes with noise masker level. (3) Relationships to behaviour: Neural tracking TRF100 was related to self-reported demand and conversations understood, while superior pariental alpha power was only related to self-reported demand.

### 4.1 Self-reported behavioural scores change concurrently with background babble noise levels, and is offset by the presence of visual cues

In the current study, the Effort subscale of the NASA-TLX was subsumed under the Mental Demand subscale to coincide with the definition of mental effort as “the deliberate allocation of mental resources to overcome obstacles in goal pursuit when carrying out a task” according to FUEL (Pichora-Fuller et al., 2016). In line with previous literature on listening effort and speech intelligibility, the current study found that self-reported mental demand ratings of CI users attending to audiovisual speech stimuli significantly increased concurrently with background noise levels, supported by similar decreases in perceived percentage of words and conversation understood. Increasing background noise has been shown to require more self-perceived listening effort as indicated by significantly increasing mean NASA-TLX scores of NH listeners attending to conversations and monologues (Peng and Wang, 2019). Accordingly, mean NASA-TLX scores of NH listeners also increases as the spectral resolution of speech (and thereby speech intelligibility) decreased in CI simulations (Pals et al., 2013).

Interestingly, mental demand ratings in the Audio-Only condition with low background noise levels (SNR+15 dB) were comparable to the demand ratings when attending to audiovisual stimuli in the High-Noise condition (SNR +5 dB). Additionally, while the perceived percentage of words understood significantly differed between the HN and AO conditions, the percentage of conversation understood did not. When looking specifically at the percentage of words and conversation understood, participants on average reported a ~20% decrease in understood words in the High-Noise condition compared to the Audio-Only condition. These results indicate that although CI users may perceive fewer words in a SNR +5 dB listening condition with visual cues, their speech perception and comprehension ability is at a similar level to that of listening without visuals in a lower noise condition. Visual speech cues have been shown to improve the ability of CI users to recognize speech (Dorman et al., 2016). The benefit of visual speech cues on speech intelligibility therefore possibly reduces the mental demand required for CI users to attend to speech in noise. However, prior research on the effect of visual speech cues on subjective listening demand are divergent. For instance, individuals reported more difficulty in understanding audiovisual speech stimuli in which the talker wore a non-transparent face mask, compared to when no face mask was worn (Yi et al., 2021). Fraser et al. (2010) observed that NH listeners were more accurate and reported less subjective effort during the audiovisual speech recognition task compared to the audio-only task in equally noisy conditions. In contrast, Brown and Strand (2019) reported that listening effort towards speech in NH listeners did not differ due to the presence of visual facial cues during a dual-task paradigm in hard listening conditions (SNR +5 dB). Reaction time was instead significantly slower in the audiovisual condition compared to the audio-only condition in easy listening conditions at SNR +10 dB (Brown and Strand, 2019).

The increase in subjective demand in the Audio-Only condition may be due to the degree to which visual speech cues influences speech perception for CI users, compared to NH listeners. CI users have been shown to gravitate towards and rely more on visual speech cues, especially in situations with poor speech intelligibility (e.g. Mastrantuono et al., 2017; Wang et al., 2020). In fact, individuals with hearing loss have been previously observed to be innately biased towards visual speech cues compared to NH listeners (Stropahl et al., 2017). Furthermore, individuals with hearing loss and CI users display cross-modal recruitment of auditory regions by visual stimuli (e.g. Anderson et al., 2017; Nishimura et al., 1999; Winn et al., 2013). Anderson et al. (2017) has shown that the post-operative increase in cross-modal superior temporal cortex activation due to visual speech cues was positively correlated with increases in speech understanding. Thus, it is possible that while CI users demonstrate greater listening effort compared to NH listeners during everyday listening, CI users also receive greater benefit from visual speech cues in the reduction of listening demand.

### 4.2 Influence of background noise and visual cues on TRF components

The current study produced speech tracking waveforms with components that have a similar morphology to potentials commonly identified as part of the cortical auditory evoked potential (i.e., N1, P2, N2). Despite the similarities between the polarity and latencies of the TRF_100_, TRF_200_, and TRF350 components to the N100, P200, and N200 event-related potentials respectively, we will not refer to these responses as the classic event-related-potentials due to differences in response characteristics as described in previous literature (Broderick et al., 2018; Reetzke et al., 2021).Recent evidence suggests that TRFs are sensitive to stimulus properties, and are more representative of low-level speech encoding (Broderick et al., 2022; Prinsloo and Lalor, 2022). Accordingly, weaker speech tracking would be indicative of either a deficit in the basic neural processing of speech acoustics, or poor encoding of low-level speech features (Broderick et al., 2022; Prinsloo and Lalor, 2022).

#### 4.2.1 No effects of increasing background noise levels on TRF component amplitude during audiovisual listening in CI users

Previous work with NH listeners has shown that background noise and attention modulate the negative early component of speech tracking around 100 ms, corresponding with our findings (Ding and Simon, 2012; Petersen et al., 2017; Verschueren et al., 2020). Petersen et al. (2017) using cross-correlations to measure neural speech tracking showed that the negative component between 124 to 160 ms time lag interval increased in magnitude for lower background noise levels. Similarly, Verschueren et al. (2020) found that as SNR decreased, the peak amplitude of the negative component within 0 to 150 ms decreased as well, even if speech intelligibility remained comparable. Increasing background noise has also been shown to also increase the magnitude of a positive speech tracking component around 200 ms time lag in individuals with hearing loss (Petersen et al., 2017). While TRF_100_ and TRF_350_ peak amplitude appeared to parametrically decrease with SNR in the current study, the difference in peak amplitude between listening conditions was non-significant. TRF_200_ peak amplitude appeared to decrease as SNR increased, but the difference did not reach significance either.

Various differences between CI users and NH listeners may explain the lack of difference in TRF component amplitudes despite the increase in background noise. Concerning the attentional modulation of speech tracking, hearing loss has been shown to negatively affect attentional modulation. Greater degrees of hearing loss were associated with changes in the tracking of the ignored speech stream, leading to greater similarities between the neural tracking of attended and ignored speech streams (Petersen et al., 2017). The lack of amplitude differences could also be due to cortical impairments involved in early-stage speech separation which has been seen in bilateral CI users (Paul et al., 2020). Interestingly, TRF_200_ components were not significantly different from the babble noise TRF waveform in all listening conditions in contrast to the TRF_100_ and TRF_350_ components compared to previous studies with NH listeners (e.g. Broderick et al., 2018; Ding & Simon, 2012). This perhaps is an indicator that TRF components represent different cognitive processes involved in neural speech tracking and attention, and that specific processes reflected in the TRF_200_ component are impaired in CI users.

Recently, Paul et al., (2020) reported that later negative speech tracking components are more representative of listening demand for people with hearing loss, due to poor early differentiation of speech. Bilateral CI users have also demonstrated stronger speech separation as reflected in later speech tracking differences around ~250 ms between attended and ignored speech in a concurrent digit-stream task, while speech tracking towards the attended speech stream was stronger in the early component around 150 ms for NH listeners (Paul et al., 2020). Although CI users in the current study demonstrated enhanced talker differentiation, the later negative component initially appeared to be enhanced for stimuli TRFs, contrary to the stronger late cortical representation of the ignored speech stream described by Paul et al., (2020). Furthermore, the mean late negative component seen in the present study peaked 100 ms later at ~350 ms, possibly as a result of using continuous speech stimuli instead of number sequences since TRF latency has been linked to stimuli prosody (Verschueren et al., 2020). Additionally, both TRF_100_ and TRF_350_ were significantly different from the babble TRF for most listening conditions, suggesting both early and late cortical speech differentiation for low and moderate noise conditions. Late cortical differentiation may therefore be more sensitive to background noise levels, as the TRF_350_ component was not statistically significantly different from the babble noise TRF in the High-Noise condition.

#### 4.2.2 Speech tracking components are not sensitive to audiovisual manipulations in low background noise scenarios

Despite the reported benefits of visual speech cues for speech perception in noise, speech tracking component amplitudes did not differ between the Low-Noise and Audio-Only conditions. The lack of difference is potentially explained by a ceiling effect regarding speech tracking. As the SNR in both audiovisual Low-Noise and Audio-Only condition was relatively high (+15 dB), participants may have been able to track the attended speech stream at the same level of performance for both conditions, despite the increased subjective listening demand in the Audio-Only condition. The benefits of visual cues then present itself in listening situations with lower SNRs, when CI users cannot adequately rely on acoustic speech cues to identify and maintain separate speech streams.

However, previous research also suggest that in audiovisual conditions, CI users favour a top-down attentional modulation process when confronted with incongruent visual information, compared to NH listeners that adopt a more bottom-up process (Song et al., 2015). Since the CI users of the current study were not instructed to close their eyes, the irrelevant visual environment in the Audio-Only condition could influence the attentional modulation process by serving as a distractor.

#### 4.3 Early speech tracking component amplitudes are correlated with self-reported demand and percentage of conversation understood, but not the percentage of words understood

The current study found that the strength of early TRF components explained mental demand ratings; more positive (i.e., weaker) TRF_100_ amplitudes are associated with increased perceived listening demand, regardless of background and audiovisual manipulations. Similarly, weaker TRF_100_ amplitudes are correlated with a decreased perceived percentage of conversation understood. The non-significant relationship observed for TRF_200_ and TRF_350_ components has also been reported in previous literature. Müller et al. (2019) measured listening effort both subjectively through self-reports and objectively using cross-correlations between the speech envelope and EEG signals, and found no significant correlations between listening effort and later component amplitudes in NH listeners.

The TRF_100_ did not significantly change with background and may be an indication of a modulatory shift of reliance from bottom-up auditory processing to top-down visual context cues for speech comprehension in increasingly non-optimal listening conditions. Increasing listening demand will force CI users to compensate by relying on visual speech cues, which may not fully recoup the effects of degradation in auditory information. The current study also found no significant correlation between AzBio scores and self-perceived percentage of conversation understood, similar to the weak correlation between clinical speech perception tests and subjective QoL outcomes seen in previous studies (e.g., McRackan et al., 2018; Ramakers et al., 2017; Thompson et al., 2020). For instance, Ramakers et al. (2017) reported weak to moderate correlations between Utrecht Sentence Test with Adaptive Randomized Roving sentences performance and the Speech, Spatial and Qualities of Hearing Scale (SSQ) (Gatehouse and Noble, 2004) Speech scale (r = −0.36, p = 0.0429) and Nijmegen Cochlear Implant Questionnaire (NCIQ) (Hinderink et al., 2000) Advanced Sound Perception Domain (r = −0.47, p = 0.0214). Similarly, Thompson et al. (2020) reported that AzBio in noise (SNR 0 dB) scores do not correlate with the Speech-in-Speech Context subscale within the SSQ Speech scale. In turn, a meta-analysis by McRackan et al. (2018) pooled correlation values from studies utilizing tests such as the SSQ and the NCIQ. The pooled correlation values ranged from negligible to low for sentence recognition in quiet (r = 0.219 [0.118 - 0.316]) and in noise (r = 0.238 [−0.054 - 0.493]) (McRackan et al., 2018). These results again highlight the complex role of visual cues during listening, as clinical hearing tests are strictly auditory in nature. Alternatively, the concurrent increase in TRF_100_ and demand ratings may represent a potential ceiling effect related to speech intelligibility (e.g. Drennan & Lalor, 2019; Verschueren et al., 2020). Whether due to low background noise or information from visual cues, participants may not have had significant issues understanding the stimuli for the neuromodulatory effects on speech tracking to significantly presents itself, despite their perception of decreased understanding.

### 4.4 Influence of background noise and visual cues on alpha power

Alpha power is theorized to increase in neural regions involved in processing information irrelevant to the task at hand (Strauß et al., 2014), possibly as a reflection of the inhibition of distracting cortical networks (Jensen and Mazaheri, 2010; Petersen et al., 2015). Changes in alpha power during listening differ depending on cortical regions of interest as well as how the sounds are presented analyzed (i.e., long continuous speech sentences or short duration sounds in event-related paradigms, e.g., see Hauswald et al., 2020). Previous studies have demonstrated that alpha power increases concurrently with task difficulty (Wöstmann et al., 2017), with increases in alpha power at the IFG in tandem with increases in listening effort (Dimitrijevic et al., 2019). Parietal alpha power was also found to increase as listening difficulty and attention increases due to increasing background noise or vocoded speech (Dimitrijevic et al., 2017; Wöstmann et al., 2017). In direct contrast, parietal and frontal alpha power has been shown to decrease as the difficulty of the listening condition increases, whether due to increasing background noise (Seifi Ala et al., 2020), decreasing target stimuli intelligibility (Fiedler et al., 2021), or vocoded speech (Hauswald et al., 2020).

Our results partially conflict with previous research, as alpha power itself at the superior-parietal and left IFG ROIs did not significantly change as an effect of increasing background noise or the presence of visual cues. One interpretation is that the effects of increasing background noise are potentially attenuated by the benefit that visual cues confer for attending to speech, reducing the required resources needed to suppress irrelevant background information. The modulation of parietal alpha power by SNR has been demonstrated for purely auditory tasks (e.g., Dimitrijevic et al., 2017; Wöstmann et al., 2017; Seifi Ala et al., 2020). The lack of difference in alpha power for the audiovisual listening conditions may therefore be an indicator of audiovisual advantage during sustained listening. At low noise levels in the Audio-Only and Low-Noise condition, background noise affects subjective listening demand, but possibly not enough to warrant neural differences in the suppression of background noise. Superior-parietal alpha power also did not differ between the side ipsilateral and contralateral to the CI side used during the listening task. While increased alpha power has been reported for the ipsilateral hemisphere to the attended stimuli (e.g. Foxe & Snyder, 2011; Paul et al., 2020; Thorpe et al., 2012) where the listening tasks involved spatial attention and the suppression of irrelevant stimuli located on the contralateral side. The target stimuli for the current study were located directly in front of the participants and did not change location. Thus, no lateralization of alpha power was expected. Stimulus material may also influence how alpha power changes as a result of degrading speech clarity. Hauswald et al. (2020) found that when using continuous stimuli parietal alpha power decreased as speech degradation increased, in contrast to studies that used short stimuli such as words or digits (Obleser and Weisz, 2012; Wostmann et al., 2015; Dimitrijevic et al., 2019). The combination of stimulus choice, the benefit of visual cues, and low background noise in some conditions may then have contributed to masking the effect of decreasing alpha power due to decreased speech comprehension across all listening conditions. Alternatively, alpha power as measured in this study may not relate to the conditions of our task design.

Regarding listening demand, a decrease in superior-parietal alpha power was associated with an increase in demand scores, but only in the audiovisual High-Noise condition. Continuous listening in an environment with high background noise would be especially difficult for individuals with hearing difficulty, perhaps to a point where CI users begin to favour visual speech cues over auditory cues to compensates for decreased speech intelligibility. Here, alpha power may represent a release from inhibition (Jensen and Mazaheri, 2010) regarding multisensory integration processes that occur in the parietal region (e.g. Misselhorn et al., 2019; Rouger et al., 2008), similar to the inverted-U pattern displayed for alpha power in high effort listening scenarios (Hauswald et al., 2020; Paul et al., 2021; Ryan et al., 2022). Any association between alpha power and demand scores for conditions with lower background noise may be complicated by CI users’ bias towards visual cues (Stropahl et al., 2017), in that CI users overestimate the benefit that visual cues confer on speech intelligibility, and thereby inherently rely more on visual cues despite the audibility of the speech stimuli. However, it is difficult to determine if this bias is purely subjective based on alpha power alone, due to the cross-modal plasticity observed in individuals with hearing loss (Lazard et al., 2010; Anderson et al., 2017) as well as the aforementioned multisensory integration. Since no visual speech cues were present in the Audio-Only condition, audiovisual integration of visual and auditory speech cues was not necessary, which would lead to the suppression of parietal multisensory integration.

It is also possible that there was not enough statistical power to detect the effects of background noise and audiovisual manipulations on TRF components and alpha power, as reflected in the variability seen in the current study sample. Some factors to be considered are the type of CI user, CI device limitations, and individual differences in effort and attention. The present study contains unilateral, bimodal, and bilateral CI users who may have different subjective listening experiences, and thus different effort and demand requirements. Bilateral CI users have reported a lower degree of listening effort compared to unilateral CI users (Hughes and Galvin, 2013; Schnabl et al., 2015; Kocak Erdem and Ciprut, 2019). Furthermore, the neural response of CI users may also have been limited by device factors such as CI distortions to the speech envelope, and thus leading to no detectable changes in TRF amplitude that corresponded with listening effort and attention. Poor encoding of the speech stimuli would thereby lead to an increase in mental demand when interpreting speech signals, in turn resulting in poor comprehension. Differences between the effort required for a task (i.e., listening effort) and the difficulty of the task itself (i.e., mental demand) may be reflected in the objective and subjective measures respectively, and thus influencing both TRF components and alpha power. If CI users did not attempt to change their listening effort to match the difficulty/demand of the task throughout all listening conditions, then the differences in both TRF amplitude and alpha power related to listening demand between conditions would be minimized, and thus not detected due to having a small effect size. As the participants were only briefly assessed on stimuli content to gauge attention, passive listening may also have taken place during the task if lapses in attention occurred. Factors beyond attention also influence both speech tracking and listening effort. Even though attentional processes affect listening effort, motivation and fatigue are also confounding factors for both concepts of listening effort and listening demand. Motivation has been found to generally increase subjective listening effort and fatigue when attending to speech in noise, in addition to influencing strategies for coping with effortful tasks (Picou and Ricketts, 2014).

### 4.5 Conclusions

Increasing background noise was detrimental to subjective mental demand when listening in noise, while visual speech cues appeared to decrease listening demand in low background noise levels. Unlike early speech tracking components, later stimuli TRF component amplitudes were not significantly different from babble noise TRFs, implicating potential difficulties in separating speech from the background. The significant relationship between early speech tracking components, alpha power, and subjective listening demand suggests that perhaps visual cues play a more complex role for CI users in higher-noise conditions. The effects of listening condition on mental demand also may be overestimated due to biases in individual perception. Relative alpha power however did not appear to change accordingly with background noise in contrast to previous literature. Nevertheless, factors such as study design, motivation, and fatigue may confound the results of neural measures for individuals with CIs compared to NH listeners, and thus warrant further investigation.

Overall, the results of this study suggest that naturalistic multi-talker scenarios such as everyday movie stimuli, can be used to study listening demand in CI users. Neural tracking of continuous naturalistic speech in noise was successfully measured in CI users using TRFs. Meaningful information about the cochlear implant listening experience and the influence of auditory environmental factors can be extracted from brain responses using continuous “ecological” stimuli such as a normal conversation. These findings contribute to the development of naturalistic objective measures of listening effort that may provide more insight into the mental effort and fatigue of CI users in everyday listening. The method of speech tracking estimation used in the current study also does not require an active response from participants, and thus has the potential to be integrated in assessments for individuals that cannot respond to conventional task-based measures. This approach may be especially useful in pediatric populations where behavioural measures of demand and long EEG recordings are difficult to perform.

## 7 Author Contributions

BX performed data analyses, interpretation, and wrote the manuscript as part of a Master of Science degree thesis. The thesis was subsequently adapted for publication. BTP and AD contributed to the conceptualization of the work, provided supervision, data analysis, and revised the manuscript before publication.

## 8 Acknowledgments

This research article was adapted from a dissertation as part of a Master of Science degree at the Institute of Medical Science at University of Toronto.

## 9 Data Availability Statement

The datasets for this study are available on request to the corresponding author.

